# Visualization of SpoVAEa protein dynamics in dormant spores of *Bacillus cereus* and dynamic changes in their germinosomes and SpoVAEa during germination

**DOI:** 10.1101/2022.01.10.475594

**Authors:** Yan Wang, Norbert O. E. Vischer, Demi Wekking, Alessandra Boggian, Peter Setlow, Stanley Brul

## Abstract

*Bacillus cereus* spores, like most *Bacillus* spores, can survive for years, germinate when their surroundings become suitable, and spore germination proteins play an important role in the initiation of germination. Because germinated spores lose dormant spores’ extreme resistance, information on the function of germination proteins could be useful in developing new strategies to control *B. cereus* spores. Prior work has shown that: i) the channel protein SpoVAEa exhibits high frequency movement in the outer leaflet of the inner membrane (IM) in dormant spores of *B. subtilis*; ii) the formation of the foci termed the germinosome between two germination proteins, the germinant receptor GerR and the scaffold protein GerD, in developing spores of *B. cereus* is slower than foci formation by GerR and GerD individually. However, the dynamics of movement of SpoVAEa in *B. cereus* spores, and the behaviour of the germinosome in *B. cereus* spore germination are unclear. In this study, we found that SpoVAEa fluorescent foci in dormant spores of *B. cereus* move on the IM, but slower than in *B. subtilis* spores, and likely colocalize transiently with GerD-mScarlet-I in the germinosome. Our results further indicate that: i) expression of GerR-SGFP2 and SpoVAEa-SGFP2 with GerD-mScarlet-I from a plasmid leads to more heterogeneity and lower efficiency of spore germination in *B. cereus*; and ii) germinosome foci observed by Fluorescence Resonance Energy Transfer (FRET) between GerR-SGFP2 and GerD-mScarlet-I b can be lost soon after the spore phase transition. However this is not always the case, as some GerR-SGFP2 foci and GerD-mScarlet-I foci continued to exist, colocalize, and even show a weak FRET signal. These data highlight the heterogeneous behaviour of spore germination protein complexes and indicate that some complexes may persist well beyond the initiation of germination.

**IMPORTANCE:** *Bacillus cereus* is commonly present in soil and does harm to humans via contaminated food. In this study, we used *B*. *cereus* spores to investigate the movement of the spore-specific channel protein SpoVAEa, the interaction between SpoVAEa and the germinosome scaffold protein GerD, as well as the dynamics of a number of germination proteins in spore germination. Our results expand upon observations of the interactions between specific *B. cereus* spore germination proteins, in particular the GerR germinant receptor A, B and C subunits and GerD, as well as between SpoVAEa and GerD. The approaches used in this work could also be used to examine the interactions between GerD and SpoVAEa and other germination proteins in spores of other *Bacillus* species.

## INTRODUCTION

*Bacillus cereus* is a Gram-positive, rod-shaped, spore forming bacterium found in soil. The vegetative cells of *B. cereus* can form endospores under harsh environmental conditions, and spores are very resistant and capable of surviving for years due to spore-specific features (1). These spore-specific properties also lead to major challenges to food safety once *B. cereus* contaminates foods, for example dairy products, rice, and chilled foods (2, 3). The major specific structural features of spores include the spore core containing chromosomal DNA, the inner membrane (IM) where germinant receptors (GRs) are located along with the germinosome scaffold protein GerD and SpoVA protein channels for the major core small molecule, a 1:1 chelate of Ca^2+^ and dipicolinic acid (CaDPA), the germ cell wall with a thin peptidoglycan layer, the cortex with a thick peptidoglycan layer, the outer membrane, the proteinaceous coat, and finally the exosporium in some *Bacillus* species, including *B. cereus* (4).

Dormant spores can initiate germination and outgrow into vegetative cells when GRs sense nutrients in the environment, such as amino acids, inosine, and sugars. Additionally, previous work indicates that GerD acts as a scaffold protein in localizing GRs in the *B. subtilis* spore IM in a complex termed a germinosome (5, 6), and the *B. cereus* GerR GR has also been shown to interact with GerD in the dormant spore IM in germinosomes (7). Another important group of spore germination proteins are the SpoVA proteins encoded by the *spoVA* operon which constitute a CaDPA channel in the IM of bacterial spores. The SpoVA proteins in spores of *B. subtilis* are SpoVAA, SpoVAB, SpoVAC, SpoVAD, SpoVAEb, SpoVAEa, and SpoVAF all encoded in one operon and expressed only in developing spores (8). The SpoVA proteins in spores of *B. cereus* are SpoVAA (BC_4070), SpoVAB (BC_4069), SpoVAC (BC_4068), SpoVAD (BC_4067), SpoVAEb (BC_4066), SpoVAEa (BC_4065) and SpoVAF (BC_4064) encoded in one operon, and SpoVAC (BC_5147), SpoVAD (BC_5148), and SpoVAEb (BC_5149) in another operon. Previous work suggests that SpoVAEa of *B. subtilis* spores, a soluble protein on the outer surface of the spore IM, may well play a role in the communication with the germinant binding GRs (9, 10). It is thus possible that SpoVAEa of *B. subtilis* spores could stimulate the opening of the SpoVA channel, thereby allowing CaDPA release (9, 10). Recent work in our lab has shown that in *B. subtilis* dormant spores SpoVAEa fused to GFP and expressed from the chromosome is present in only one focus which exhibits random high frequency movement on the spore IM (11). However, SpoVAEa and the germinosome scaffold GerD protein did not interact in pulldown assays using extracts from *B. subtilis* spores (Y.-Q. Li and B. Hao, unpublished data discussed in ref. (9)). Therefore, strong interaction between these proteins was not observed, although we cannot exclude the existence of transient interaction between SpoVAEa and GerD. There is also only minimal understanding of the location and physical state of SpoVAEa in dormant spores of the food pathogen *B. cereus*, nor whether SpoVAEa and GerD proteins colocalize at least even transiently.

The spore germination process in Bacilli and Clostridia has been reviewed in the past years (12–14). Initially germinants bind to GRs, followed by large-scale release of monovalent cations and then CaDPA release via the SpoVA protein channel (15, 16). The kinetics and heterogeneity of spore germination triggered by L-alanine have been analysed giving the frequency distribution at both the population level and in individual spores of *B. cereus* strain T using phase-contrast and fluorescence microscopy (17–19). Previous work showed that the GR GerR is in the spore IM using fluorescent reporter protein fusions and the membrane dye FM 4-64, and the GerR GR is primarily responsible for L-alanine germination of *B. cereus* spores (7, 20, 21). Recent work also showed that in the IM germinosome, GerR and GerD could be visualized using fluorescent reporter proteins and this work suggested that the formation of germinosome foci was significantly slower than the formation of GerR-SGFP2 and GerD-mScarlet-I foci, and with significant heterogeneity in formation of germinosome foci (7). However, there is little information about the overall changes or behaviour of these germination proteins during *B. cereus* spore germination.

The strongly enhanced green fluorescent protein (SGFP2) and mScarlet-I have been successfully used to visualize the spore germination proteins GerR and GerD in spores of *B. cereus* ATCC 14579 when fluorescent fusion proteins were expressed from a low-copy number plasmid expression vector (7, 20). In this work, we aimed to visualize the movement of SpoVAEa with SGFP2 in dormant spores of *B. cereus* ATCC 14579 using fluorescence microscopy, and to analyze the fluorescence distribution by the changes, either up or down, in the full width at half maximum (FWHM) of the fluorescence. Additionally, the phase contrast intensity and the fluorescence changes of germinosome foci formed by GerR-SGFP2 and GerD-mScarlet-I, and SpoVAEa-SGFP2 and GerD-mScarlet-I were tracked by a time-lapse microscope equipped with phase-contrast and fluorescence analysis options. This work found that SpoVAEa-SGFP2 foci, one or multiples per spore, exhibited random movements in the IM, and often colocalized with GerD-mScarlet-I in dormant *B. cereus* spores. The results also suggested that expression of GerR and SpoVAEa proteins with GerD affected germination efficiency and led to slower and more heterogeneous spore germination. Upon addition of germinant to spores and the initiation of germination, intensities of germinosome FRET foci were lost most rapidly followed by decreases in the intensities of GerR-SGFP2 and then GerD-mScarlet-I foci. Yet some GerR-SGFP2 foci and GerD-mScarlet-I foci continued to exist and remained colocalized even after the spores’ transitioned from phase bright to dark, suggesting that germinosome-like complexes may persist beyond completion of germination. Loss of SpoVAEa-SGFP2 fluorescence intensity also occurred beginning upon spores’ transition from phase bright to phase dark.

## RESULTS

### Movement of SpoVAEa foci in dormant spore of *B. cereus*

Previous studies showed that fluorescence full width at half maximum (FWHM) can be used to quantitate fluorescence distribution of spore proteins (11). In this study, we used dormant spores of *B. cereus* strain 014 expressing SpoVAEa-SGFP2 from a plasmid to observe the movement of SpoVAEa-SGFP2 foci. The percent changes of FWHM in 100 frames of an individual *B. cereus* spore over a 5 sec period were calculated as up (positive percentage) or down (negative percentage) compared to the first frame. This work found that SpoVAEa-SGFP2 indeed showed different what appeared to be random movements or flexing in *B. cereus* spores 1, 2, 3 and 4 (Fig. 1A, Fig. S1). However, the percent changes in SpoVAEa-SGFP2 FWHM in *B. subtilis* spores exhibited a wider boundary and higher frequency changes, either up or down, compared to *B. cereus* spore 2 (Fig. 1B). This result suggested that the SpoVAEa-SGFP2 foci in individual spores of *B. cereus* and *B. subtilis* both moved in the IM and thus potentially could interact with germinosome components. Of note, SpoVAEa fluorescent foci in *B. subtilis* spores redistributed at a higher frequency than those in *B. cereus* spores. This difference may be because different species with a different protein complement are being compared, and with genomic expression of the fluorescent fusion protein in *B. subtilis* versus expression from a plasmid in *B. cereus*.

**FIG 1.**
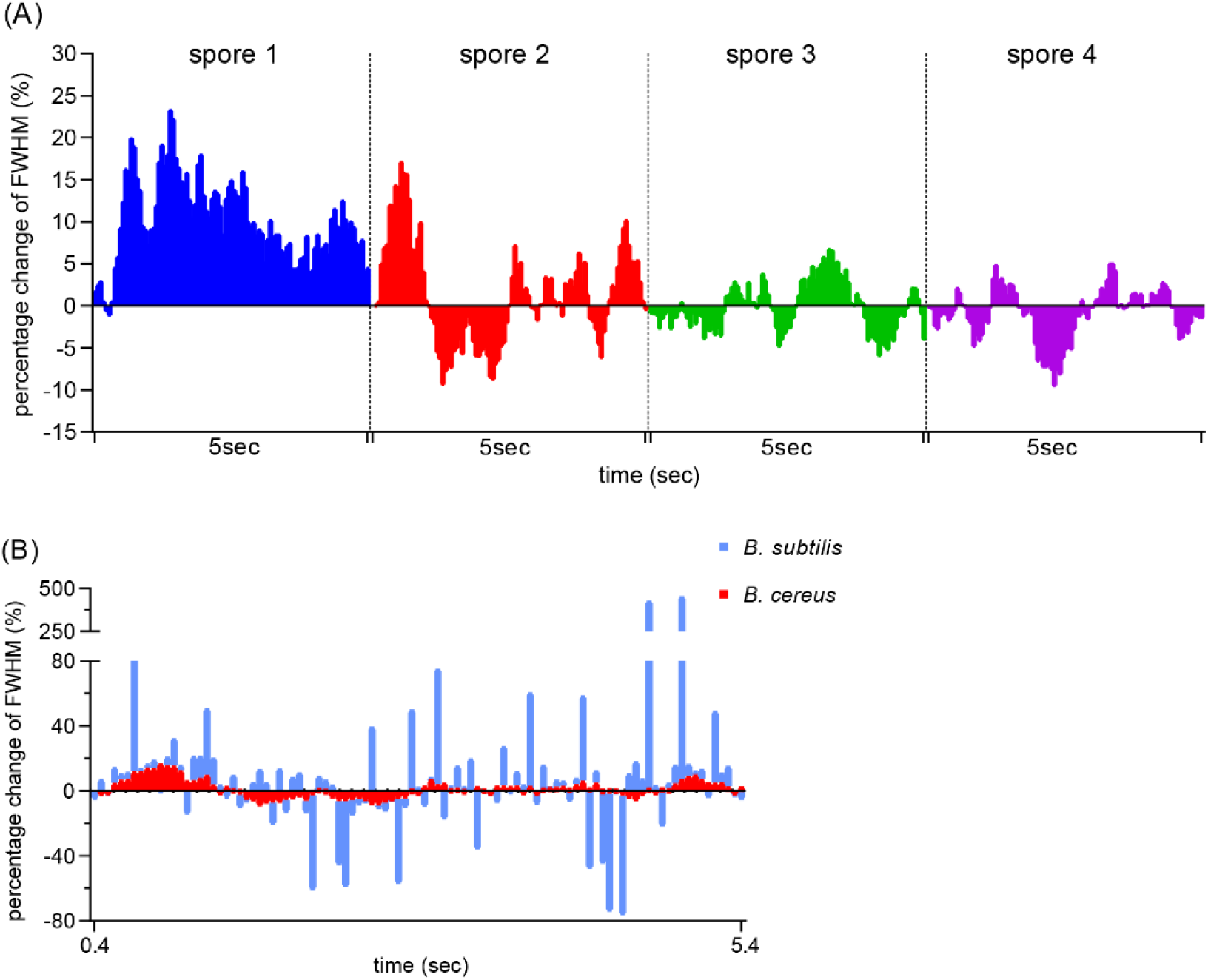
Comparison of the movements of SpoVAEa-SGFP2 foci in dormant spores of *B. cereus* and *B. subtilis.* Panel A, the percentage changes of FWHM in individual spores 1, 2, 3 and 4 of *B. cereus.* Panel B, the percentage changes of FWHM in *B. cereus* spore 2 (red squares) and a *B. subtilis* spore (blue squares). The positive and negative percentages of columns indicate the up and down of the tendency of fluorescence FWHM distribution. The montage of 100 frames (A1 to J10) of *B. cereus* spore 2 is given in Fig. S1.

### SpoVAEa-SGFP2 levels are enhanced by GerD expression in recombinant *B. cereus* spores

Our recent work showed that GerR and GerD foci are present and colocalized in germinosomes of dormant spores of *B. cereus* (20). In current work, fusion protein SpoVAEa-SGFP2 was expressed alone in spores of *B. cereus* strain 014, or expressed with GerD-mScarlet-I in spores of strain 015. The fluorescence intensities of SGFP2 in strains 014 and 015 are, as expected, both higher than those of wild type *B. cereus* spores without the recombinant proteins. However, it is notable that the fluorescence in spores of *B. cereus* strain 015 looks brighter than in spores of strain 014 (Fig. 2A). Indeed, when the fluorescence levels in the spores of recombinant strains 014 and 015 were compared, the distribution of fluorescence intensity in a spore population ranging from one to over five-fold the wild-type level was skewed to the right in *B. cereus* strain 015 spores compared to strain 014 spores (Fig. 2B). Thus, when SpoVAEa-SGFP2 and GerD-mScarlet-I were expressed from the same plasmid, GerD may contribute to the stability of the SpoVAEa, or perhaps enhance SpoVAEa expression.

**FIG 2.**
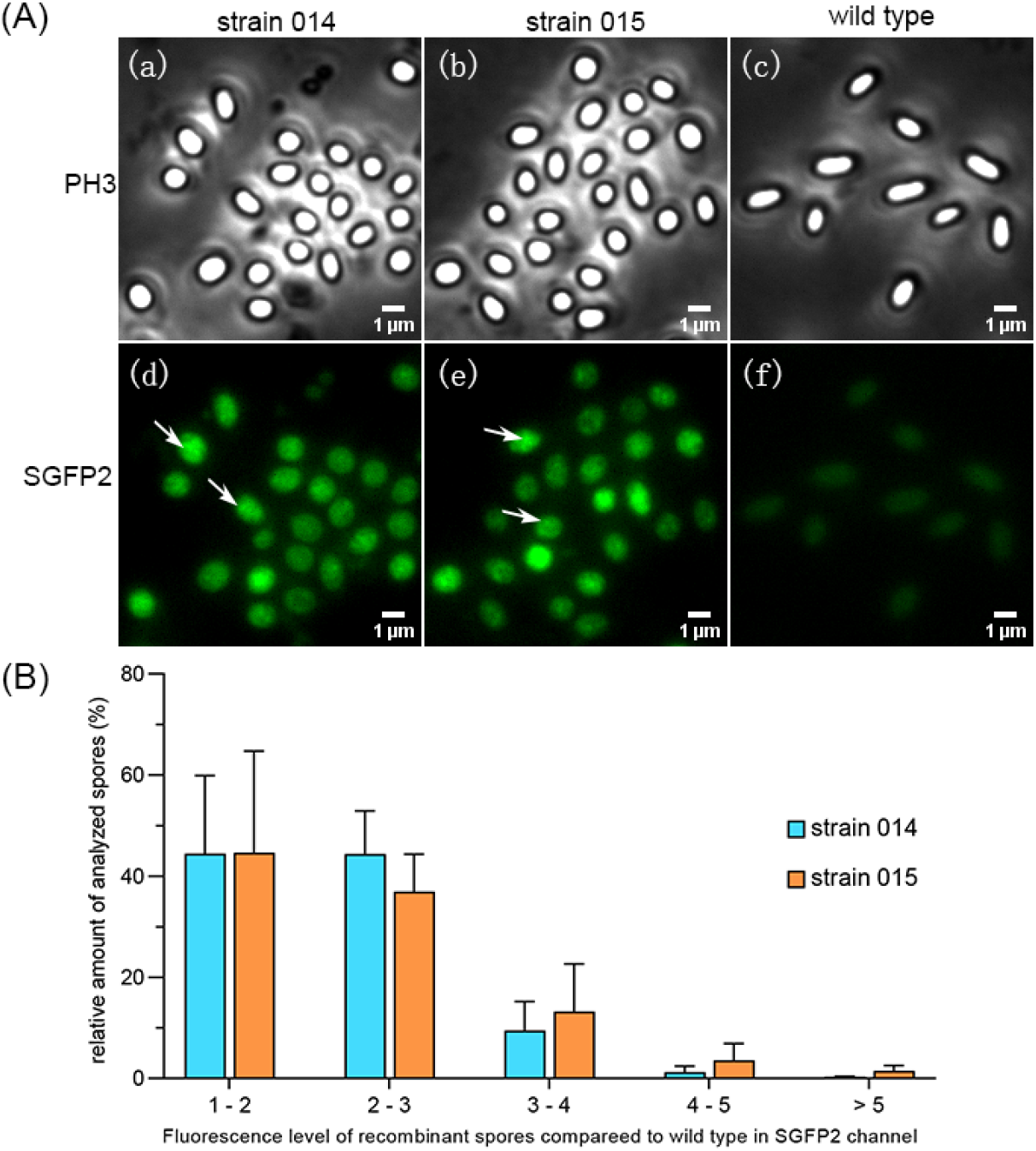
Visualization and comparison of SpoVAEa-SGFP2 fusion protein fluorescence in dormant spores of *B. cereus* strains 014 and 015. A) Dormant spores of *B. cereus* strains 014 expressing SpoVAEa-SGFP2, 015 expressing both SpoVAEa-SGFP2 and GerD-mScarlet-I and wild type were visualized: (a), (b), and (c) in the phase contrast (PH3) channel, or (d), (e), and (f) in the SGFP2 fluorescence channel. Note the multiple SpoVAE-SGFP2 foci in individual 014 and 015 spores. B) Fluorescence levels in the SGFP2 channel in spores of strains 014 and 015 compared to wild type. Data are shown as a mean with SD.

### Analysis of colocalization of SpoVAEa and GerD proteins

Given the results described above, it was important to examine possible interaction between SpoVAEa-SGFP2 and GerD-mScarlet-I in dormant *B. cereus* spores. The spectra of SGFP2 and mScarlet-I have an overlap that may produce a larger Pearson’s coefficient, which is a commonly used colocalization indicator, (7, 22, 23). To reduce effects of GerD-mScarlet-I itself, spores of *B. cereus* strain 007 expressing GerD-mScarlet-I alone were used as a control. The analysis (Fig. 3) showed that the Pearson’s coefficient of SGFP2 and mScarlet-I channels in spores of *B. cereus* strain 015 was significantly higher than those in the control. This result indicated that there is likely colocalization, albeit perhaps only transiently, between SpoVAEa-SGFP2 and GerD-mScarlet-I proteins in *B. cereus* spores.

**FIG 3.**
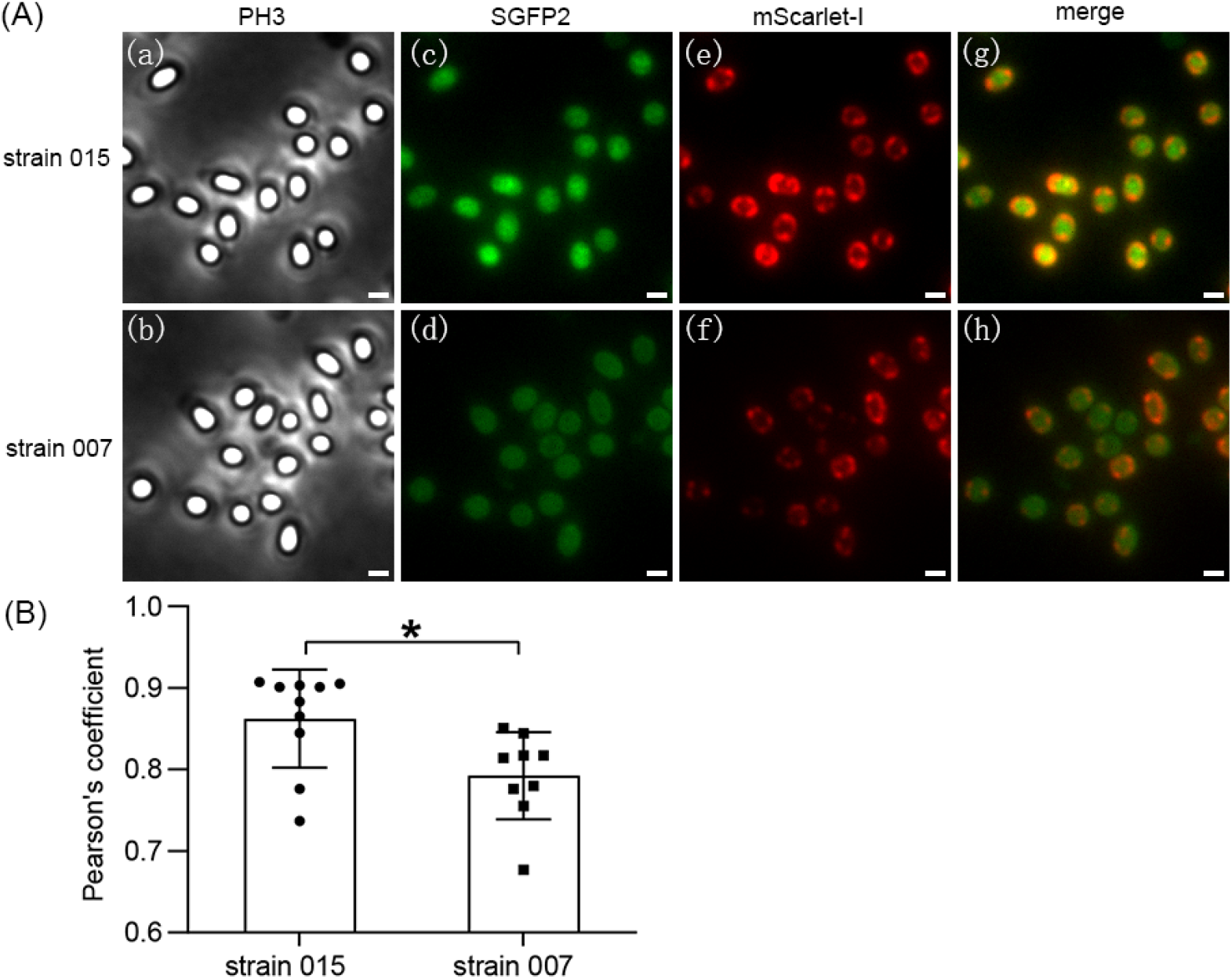
Analysis of colocalization of SpoVAEa and GerD proteins in dormant spores of *B. cereus* strain 015. Panel A, visualization of *B. cereus* strain 015 spores expressing SpoVAEa-SGFP2 and GerD-mScarlet-I and 007 spores expressing only GerD-mScarlet-I: (a) and (b) Phase-contrast channel (PH3); (c) and (d), SGFP2 channel; (e) and (f), mScarlet-I channel; (g) and (h), merged image of SGFP2 and mScarlet-I channels. The scale bar is 1 μm. Panel B, Pearson’s coefficient between SGFP2 and mScarlet-I channels. Data are shown as a mean with SD. *, *P* < 0.05.

### Expression of GerR and SpoVAEa with GerD affects *B. cereus* spore germination

In the current work, the time of initiation of germination is termed germX, which is defined as the time of the beginning of the rapid decrease in a spores’ phase contrast image intensity. When GerR-SGFP2 and SpoVAEa-SGFP2 with GerD-mScarlet-I were expressed from a plasmid in *B. cereus* strains F06 or 015, respectively, germX values exhibited greater heterogeneity than in spores of *B. cereus* strains 006 and 014 (Fig. 4, 5). In particular, with individual spores of *B. cereus* strain 007 expressing GerD-mScarlet-I alone from a plasmid, their germX values exhibited somewhat more heterogeneity than those of spores of strains 006 and 014 (Fig. 4, 5). These results indicated that expression of GerD-mScarlet-I from a plasmid in *B. cereus* can lead to more heterogeneity in spore germination.

**FIG 4.**
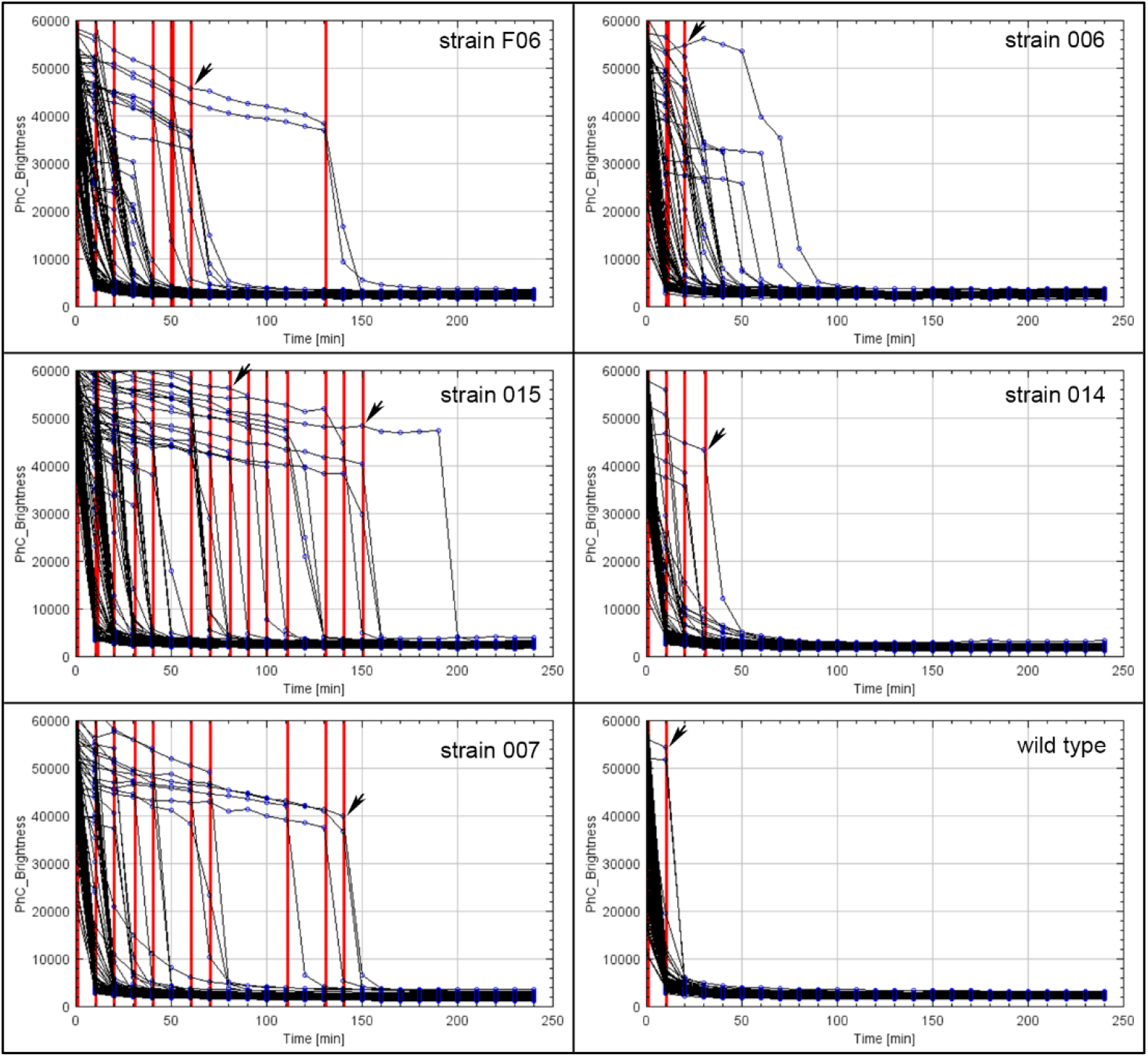
The phase plots show the germination of spores of *B. cereus* strains F06, 006, 015, 014, 007 and wild type. The red lines and black arrows indicate the time of initiation of spore germination, termed germX, for each spore. For each strain, each black line indicates the change of phase-contrast intensity in an individual spore during germination. The numbers of analyzed spores of *B. cereus* strain F06 expressing GerR-SGFP2 and GerD-mScarlet-I, strain 006 expressing GerR-SGFP2, strain 015 expressing SpoVAEa-SGFP2 and GerD-mScarlet-I, strain 014 expressing SpoVAEa-SGFP2, strain 007 expressing GerD-mScarlet-I and wild type were 108, 92, 122, 233, 107 and 265, respectively.

**FIG 5.**
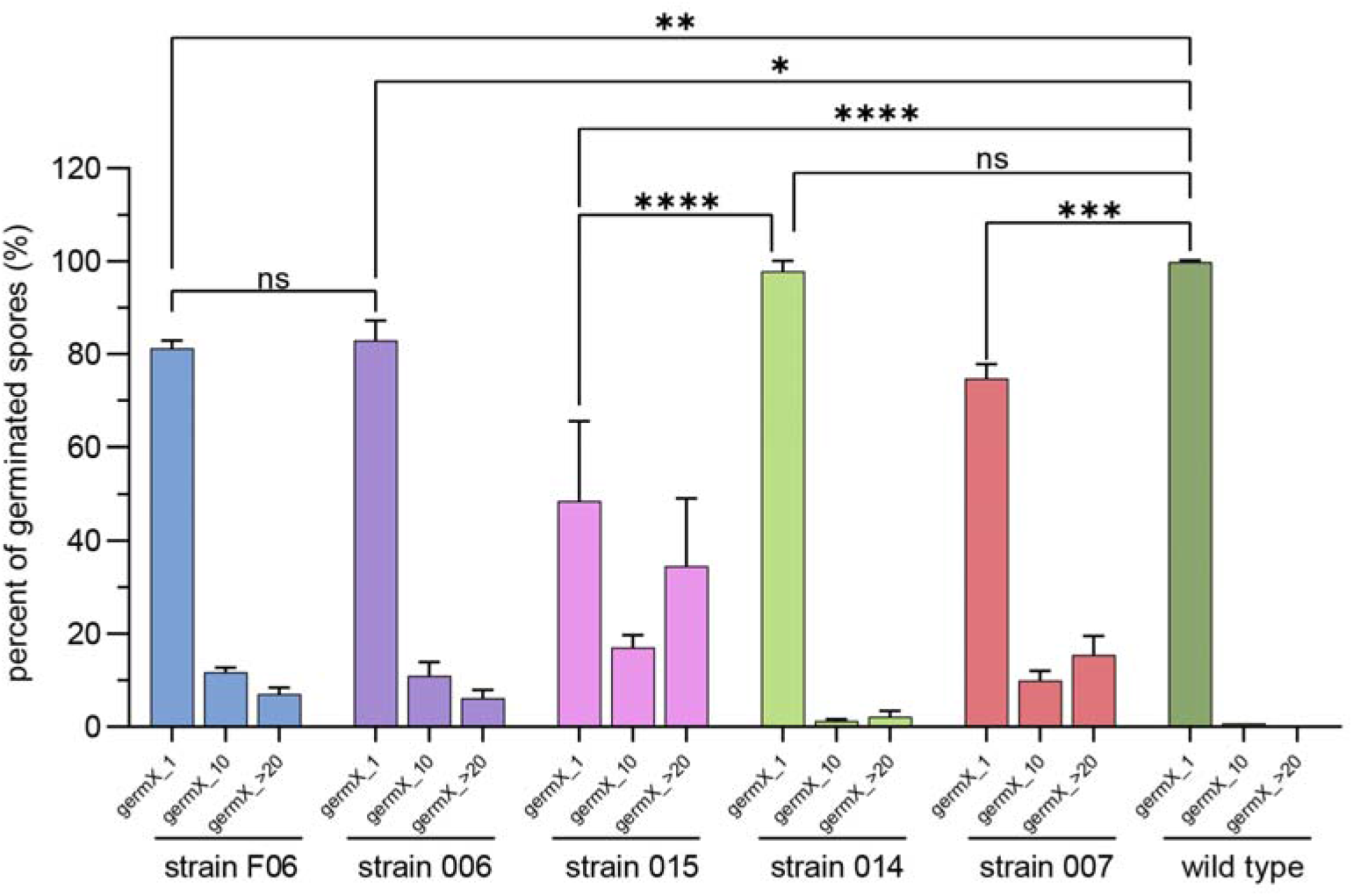
Germination of spores of *B. cereus* strain F06 expressing fusion proteins GerR-SGFP2 and GerD-mScarlet-I, strain 006 expressing fusion protein GerR-SGFP2, strain 015 expressing fusion proteins SpoVAEa-SGFP2 and GerD-mScarlet-I, strain 014 expressing SpoVAEa-SGFP2, strain 007 expressing fusion protein GerD-mScarlet-I, and wild type were analyzed based on the different germX times: germX_1, initiation of germination at ≤1 min; germX_10, initiation of germination at <10 min; germX_>20, initiation of germination >20 min. Data are shown as the mean with SD and are averages of three independent experiments. The numbers of analyzed germinated spores of different *B. cereus* strains are listed in Table S2. ns, not significant; *, *P* < 0.05; **, *P* < 0.01, ***, *P* < 0.001; ****, *P* < 0.0001.

Previous work has shown that expression of the L-alanine-responsive GerA GR controlled by the strong forespore-specific *sspB* promoter in the *B. subtilis* genome can significantly increase the rate of germination triggered by L-alanine (24). Our results showed that spores of all different *B. cereus* strains started to germinate by 1, 10 and >20 min, termed groups germX_1, germX_10, and germX_>20. Most spores were in the germX_1 group, with 81.2%, 82.9%, 48.5%, 97.7%, 74.7%, and 99.8% of spores of *B. cereus* strains F06, 006, 015, 014, 007 and wild type, respectively (Fig. 5). However, plasmid expression of fusions of GerR with or without GerD in strains F06 and 006 led to significantly lower germination efficiency compared to wild type spores, with effects in F06 spores slightly greater than in 006 spores. This suggests that increased GerR expression may not increase rates of spore germination with L-alanine. However, this only a suggestion since: i) levels of GerR-SGFP2 and GerR itself are not known in these spores; and ii) it is not known if GerR-SGFP2 can function in spore germination or might even exert a dominant negative effect on L-alanine germination although GerR-SGFP2 is certainly competent in germinosome formation. The germination results further showed that plasmid expression of SpoVAEa alone in spores of strain 014 also had no significant effect on germination efficiency compared to that of wild type spores. However, expression of GerD with SpoVAEa fusion proteins in spores of *B. cereus* strain 015 significantly (*P* < 0.0001) slowed germination compared to wild type spores or spores expressing only one of the fusion proteins (Fig. 4, 5). Notably, expression of GerR-SGFP2 alone or together with GerD-mScarlet-I in strains 006 or F06, respectively, led to a significantly lower germination efficiency compared to wild type spores (Fig. 5). However, F06 and 006 spores’ germination was not significantly faster than that of strain 007 spores that expressed the GerD fusion protein alone (data not shown).

Another important piece of information from the results in Figure 5 is that spores of strain 014 containing only SpoVAEa-SGFP2 exhibited minimal if any change in germination from that of wild type spores. The importance of this result is that recent work (25) has shown that *B. cereus* spores of strains carrying the plasmid used in the current work exhibited a significantly altered protein composition compared to that in plasmid-free spores, with many hundreds of spore proteins significantly upregulated and downregulated. Since the plasmid backbone in all constructs used in the current work is the same, the fact that spores of strain 007 and wild type spores gave almost identical germination profiles indicates that the presence of the plasmid backbone alone does not alter spore germination. Thus, effects of plasmids containing fusion proteins on spore germination kinetics seen in this work can be attributed to the fusion proteins expressed from the plasmids.

### Dynamics of germinosome behaviour upon germination triggered by L-alanine in *B. cereus* spores

Our recent study suggested that the formation of FRET foci between GerR-SGFP2 and GerD-mScarlet-I could be significantly slower than the formation of foci in the SGFP2 and mScarlet-I channels during *B. cereus* spore formation (7). In the current work, we aimed to track the dynamic changes in germinosome FRET foci upon germination triggered by L-alanine in spores of *B. cereus* strain F06 expressing GerR-SGFP2 and GerD-mScarlet-I from a plasmid. The phase-contrast channel (PH3) recorded the transition between a phase-bright individual spore to a phase-dark spore at 1 min and 10 min of the germination time in the germX_1 and germX_10 groups, respectively (Fig. 6A, Table S3). Upon the initiation of this phase transition in the germX_1 group of strain F06 spores, the intensity of germinosome FRET foci fell significantly in comparison to the beginning of the experiment (0 min), but there was no significant decrease in the intensity of GerR-SGFP2 foci (Fig. 6B, Table S3). The average NFRET values in spores of strain F06 at 10 min, 20 min, 30 min, 40 min, 50 min and 60 min of the germX_1 group were all lower than those at 0 min, but again no significance could be established for this downward trend, indicative of heterogeneous behaviour of individual spores within the population (Fig. 6C). Note that while Fig. 6A gives results for an individual spore, Figs. 6B and 6C represent population averages. When GerR-SGFP2 alone and GerD-mScarlet-I alone were expressed in spores of strains 006 and 007, respectively, the intensities of 006 spores in the SGFP2 channel in the germX_1 group at 20 min were significantly lower than at 0 min (*P* < 0.01). The intensities of 007 spores in the mScarlet-I channel of the germX_1 group at 20 min were also significantly lower than those at 0 min (*P* < 0.001) (Fig. S2, Table S4). The intensities of the FRET foci of F06 germX_10 spores at 40 min and beyond were also significantly lower in comparison with the 0 min spores. The FRET intensity drop occurred after the initiation of the rapid fall in spores’ phase contrast image intensity starting at 20 minutes after the initiation of our experiment (Fig. 6B, Table S3). In addition, remaining GerD-mScarlet-I foci became less intense in germinated spores of the germX_1 and germX_10 groups (Fig. 6B). The intensities of strain 006 spores expressing GerR-SGFP2 alone in the SGFP2 channel of the germX_10 group showed a downward trend during our experiment except from the 30 min time point. The same trend was observed for the 007 spores albeit that the drop in the intensities of the mScarlet-I channel only started 20 min after the start of our experiment (Fig. S2, Table S4). The NFRET values in spores of strain F06 at 10 min, 20 min, 30 min, 40 min, 50 min and 60 min of the group germX_10 were all lower than those at 0 min, but also here no significance could be established to the downward trend, indicative again of heterogeneous behaviour of individual spores within the population (Fig. 6C). While these results indicate that the germinosome FRET foci in spores of *B. cereus* can be lost soon after the spore phase transition, this is not always the case, as some GerR-SGFP2 foci and GerD-mScarlet-I foci continued to exist, colocalize, and even show a weak FRET signal (compare Figure 6A left- and right-hand panels). These data highlight the heterogeneous behaviour of spore germination protein complexes and indicate that some complexes may persist well beyond the initiation of germination.

**FIG 6.**
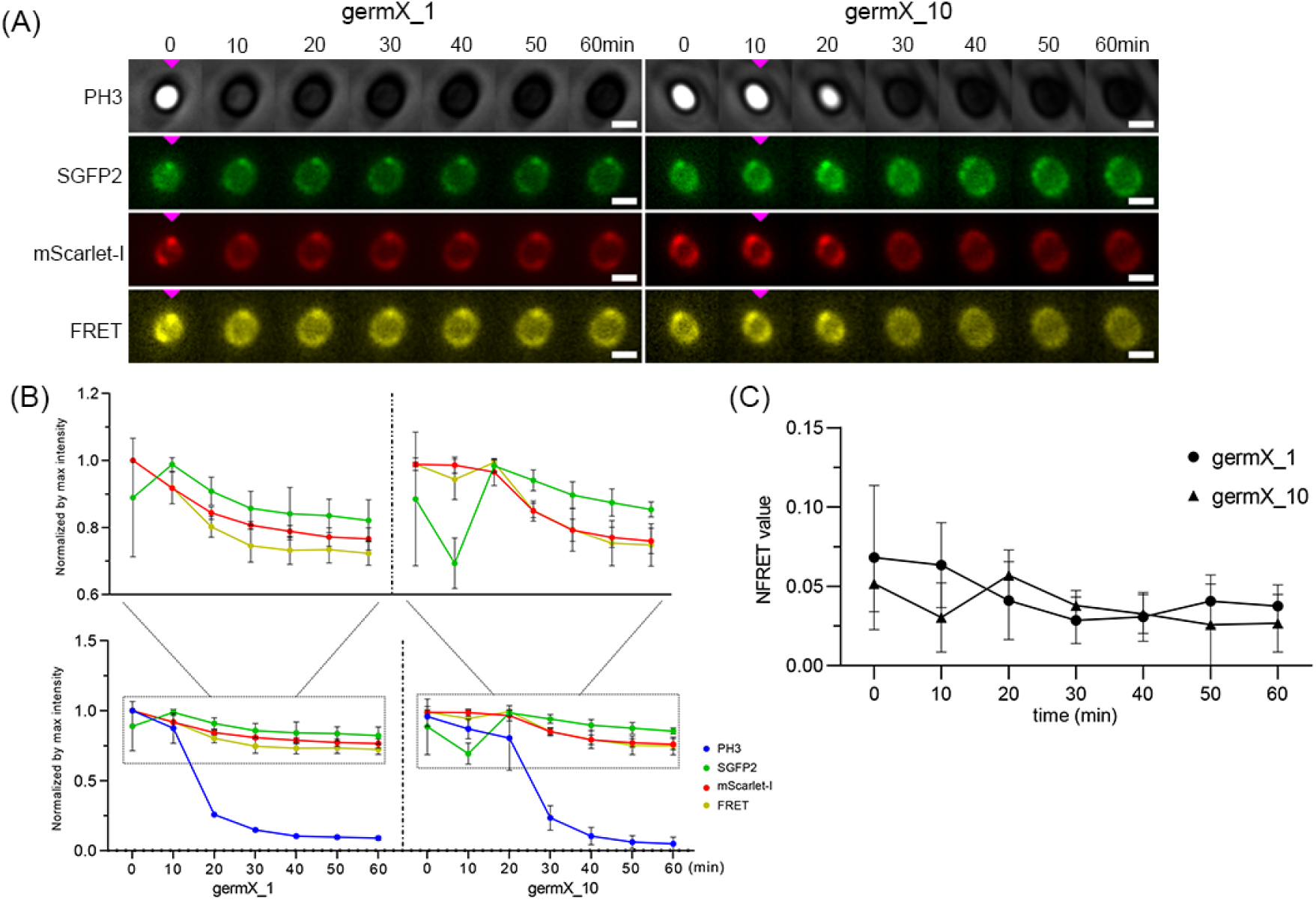
Dynamic changes in germinosome foci upon L-alanine germination of *B. cereus* strain F06 spores expressing GerR-SGFP2 and GerD-mScarlet-I. Panel A, visualization of changes in GeR-SGFP2, GerD-mScarlet-I and germinosome foci at 10 min intervals over 60 min. The left column is the phase contrast (PH3), SGFP2, mScarlet-I and FRET channels of an individual spore in the germX_1 group. The right column is phase contrast (PH3), SGFP2, mScarlet-I and FRET channels of an individual spore in the germX_10 group. The pink triangles indicate the time of initiation of germination. The scale bar is 1 μm. Panel B, the line charts of intensities in the PH3, SGFP2, mScarlet-I and FRET channels of spore populations. Left column, germX_1 group; right column, germX_10 group. Panel C, the line chart shows the dynamics of normalized FRET (NFRET) values in populations of spores of strain F06. The black circles represent the germX_1 group. The black triangles represent the germX_10 group. Data are shown as the mean with SD. The number of analysed germinated spores of *B. cereus* strain F06 is listed in Table S2. The statistical analyses between individual time points in line charts in comparison to the previous one are given in Table S3.

### Dynamics of SpoVAEa and GerD proteins during germination of *B. cereus* spores triggered by L-alanine

In this work, recombinant spores of *B. cereus* strain 015 expressing SpoVAEa-SGFP2 and GerD-mScarlet-I fusion proteins from a plasmid were used to visualize the dynamic changes of SpoVAEa and GerD upon germination initiated by L-alanine. The results showed that the phase-contrast intensity of germinated spores of strain 015 at 10 min in the germX_1 group was greatly decreased compared to that of phase-bright spores at 0 min. In the germX_10 group, the phase transition occurred after 10 min and the phase-contrast intensity of germinated spores at 20 min was decreased compared to that at 0 min (Fig. 7, Table S5). The fluorescence intensity in the SGFP2 or mScarlet-I channels of germinated spores at 10 min in the germX_1 group was decreased compared to that of phase-bright spores at 0 min, but this reduction was not significant (*P* > 0.05). Instead, when SpoVAEa-SGFP2 alone was expressed in strain 014 spores, the intensities of strain 014 spores in the SGFP2 channel of the germX_10 group at 10 min was significantly decreased compared to that at 0 min (Fig. S2, Table S4). The results showed that the SpoVAEa-SGFP2 foci were lost, and overall spore green fluorescence intensity dropped following initiation of germination. The same is true for GerD-mScarlet-I, although, in accordance with our results described in Fig. 6, some foci continued to exist beyond the phase transition albeit with lower fluorescent intensity.

**FIG 7.**
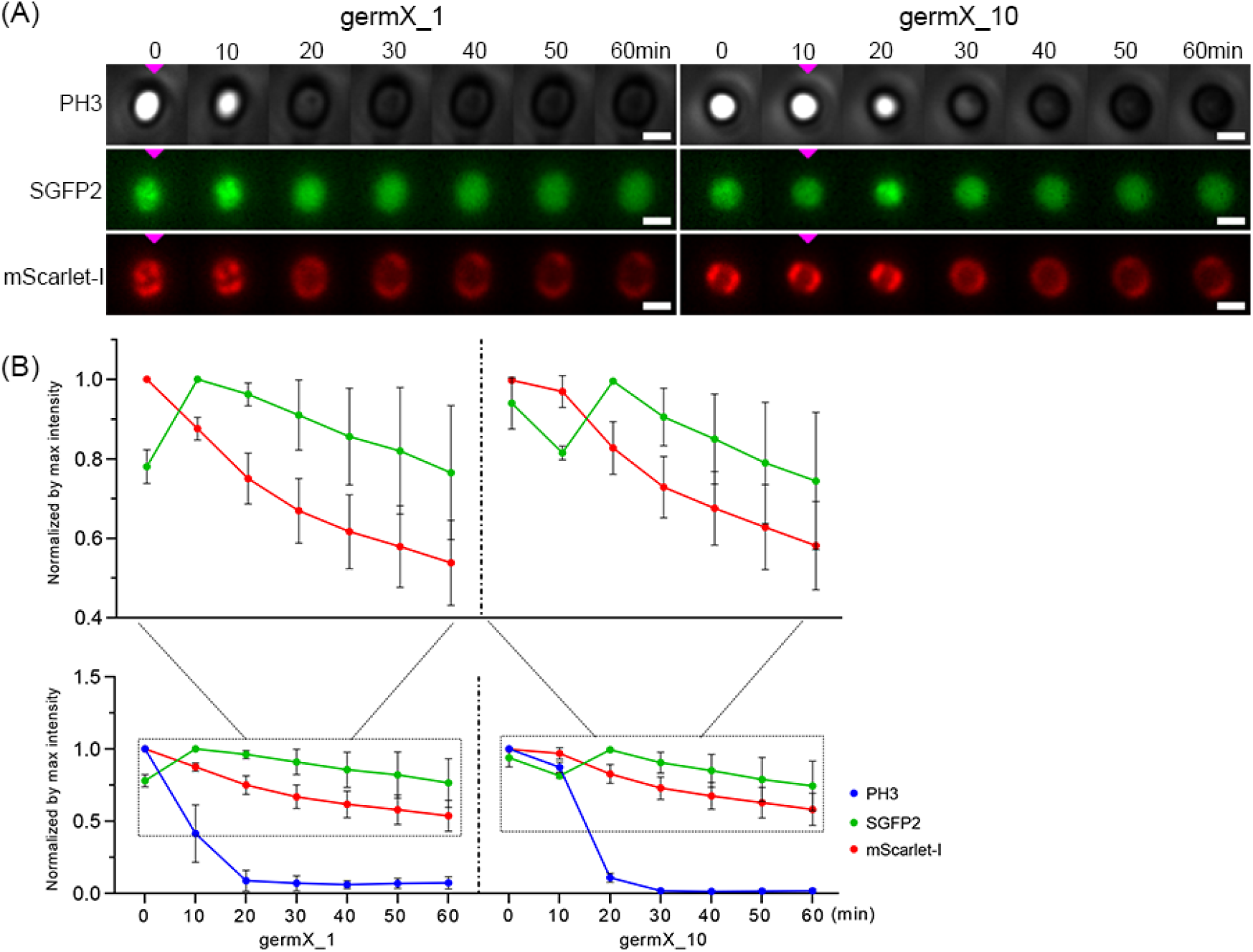
Dynamic changes in SpoVAEa-SGFP2 and GerD-mScarlet-I fluorescence intensities during L-alanine germination of *B. cereus* strain 015 spores. Panel A, visualization of changes in SpoVAEa-SGFP2 and GerD-mScarlet-I at 10 min intervals over 60 min in one spore. The left column is phase contrast (PH3), SGFP2 and mScarlet-I channels of an individual spore in the germX_1 group. The right column is PH3, SGFP2 and mScarlet-I channels of an individual spore in the germX_10 group. The pink triangles all indicate the time of initiation of germination. The scale bar is 1 μm. Panel B, the line charts of PH3, SGFP2 and mScarlet-I channels from multiple individual spores. Left column, germX_1 group; right column, germX_10 group. Data are shown as the mean with SD and represent three independent experiments. The numbers of analysed germinated spores of *B. cereus* strain 015 are listed in Table S2. The statistical analysis of the time points in line charts in comparison to the previous one is given in Table S5.

## DISCUSSION

*B. cereus* spores, like most *Bacillus* spores, have various resistance characteristics due to spore specific structures, and can restart metabolism only after spore germination has been completed. The nutrient germination of spores is initiated by germinant binding to specific GRs localized in spore’s IM, including GerR, GerK, GerG, GerL, GerQ, GerI, and GerS in spores of *B. cereus*, with GerR triggering germination with L-alanine (21, 26). In addition, SpoVAEa is a component of the IM SpoVA protein channel for CaDPA and GerD is a scaffold protein playing an important role in germinosome formation and thus spore germination in *B. subtilis* and *B. cereus* (6, 7, 9). To extend these latter observations, we have now studied the dynamic changes of the SpoVAEa protein in dormant and germinated spores of *B. cereus*, and the kinetic changes in germinosome foci during the germination process.

Based on previous observations, the expression levels of GerD and most SpoVA proteins in *B. subtilis* spores are ~10^2^-fold and 10^3^-fold higher than those of GRs, respectively (27). Fluorescence microscopy of *B. cereus* spores showed clear foci of GerD-mScarlet-I and less distinct foci of SpoVAEa-SGFP2, and a possible reason for this could be the different levels of these two proteins. However, the weaker fluorescent signal of SpoVAEa-SGFP2 than GerD-mScarlet-I in our work might also be caused by expression of only one subunit of the SpoVA complex alone, or perhaps SpoVAEa has a lower expression level than other SpoVA proteins, as was shown for SpoVAEa versus SpoVAD in *B. subtilis* spores (9).

In this study, recombinant *B. cereus* spores expressing fluorescent fusions to GerR and SpoVAEa with or without a GerD fusion protein from plasmids were used to assess the effects of these fusion proteins on germination triggered by L-alanine. Plasmid expression of GerD-mScarlet alone or with SpoVAEa-SGFP2 clearly slowed spore germination, as shown by increases in the levels of spores in the germX_10 and germX_>20 groups in the spores expressing GerD-mScarlet-I (Fig. 5; Table S2). These results indicated that plasmid expression of the GerD fusion protein has an inhibitory effect on spore germination efficiency and thus increased germination heterogeneity, consistent with previous work (17, 19). However, spores with plasmid expression of the GerR fusion alone also germinated more slowly than wild type spores, but co-expression of the GerD fusion did not slow germination any further (Fig. 5, Table S2). The reasons for the effects of the various fusion proteins on rates of spore germination are not clear. Two possible reasons are that the protein fusions are either not functional in germination or exert a dominant-negative effect on the function of the wild-type proteins. While the latter are possibilities, the GerD and GerR fusion proteins do form germinosomes (7), so retain at least some wild-type function. In addition, GerD and GR fusion proteins analogous to those used in the current work were functional in *B. subtilis* spore germination (28). However, ultimately answering this functionality question definitively will require analysis of the effects of the fusion proteins in the appropriate null nutant backgrounds. Notably, a recent study suggested that in *B. subtilis*, the B subunit of the GerA GR, GerAB, is responsible for binding the germinant L-alanine, and this is consistent with molecular dynamics analyses of L-alanine binding to GerAB (29). Our recent studies also suggest that there is a very close interaction between GerRB and GerD in the germinosome. Consequently, the reason for the inhibitory effect of plasmid GerD-mScarlet expression plus or minus SpoVAEa-SGFP2 on spore germination could be that excess GerD can occupy or occlude the L-alanine binding sites on GerRB, while plasmid expression of GerR-SGFP2 would increase GerRB subunit levels such the concomitant expression of the GerD protein fusion has minimal effects on spore germination.

Our recent results, including using FRET-based analysis, on the dynamics of germinosome formation in *B. cereus* spores suggest that the formation of foci in the FRET channel may be significantly slower than formation of the GerR-SGFP2 and GerD-mScarlet-I foci (7). To further assess germinosome dynamics, we observed the changes in germinosome foci upon germination initiated by L-alanine in *B. cereus* spores. In this experiment, the protein FRET pairs, GerR-SGFP2 and GerD-mScarlet-I, were expressed from a plasmid and driven by their native promoters during sporulation. Possibly consistent with the role of the B subunit of GerA in *B. subtilis*, GerRB may also be responsible for initiating germination with L-alanine in *B. cereus* (Fig. 6). Once the process of spore germination was initiated, our results showed that some GerD - GerR colocalization likely remains, as even though FRET positive germinosome foci were lost after the initiation of germination, some GerD and GerR complexes may continue to exist beyond this time point. Fig. 8 shows a hypothetical sequence of events that may occur during spore germination. A note of caution is warranted because the germination proteins analysed in this work were expressed from a plasmid and evidently may disrupt the dynamic balance in germination protein assembly in sporulation and germination. Importantly though, all germination proteins studied were expressed from the plasmid under the control of their respective native promoters allowing relative expression differences to be conserved, although this will likely be influenced by plasmid copy number which is variable.

**FIG 8.**
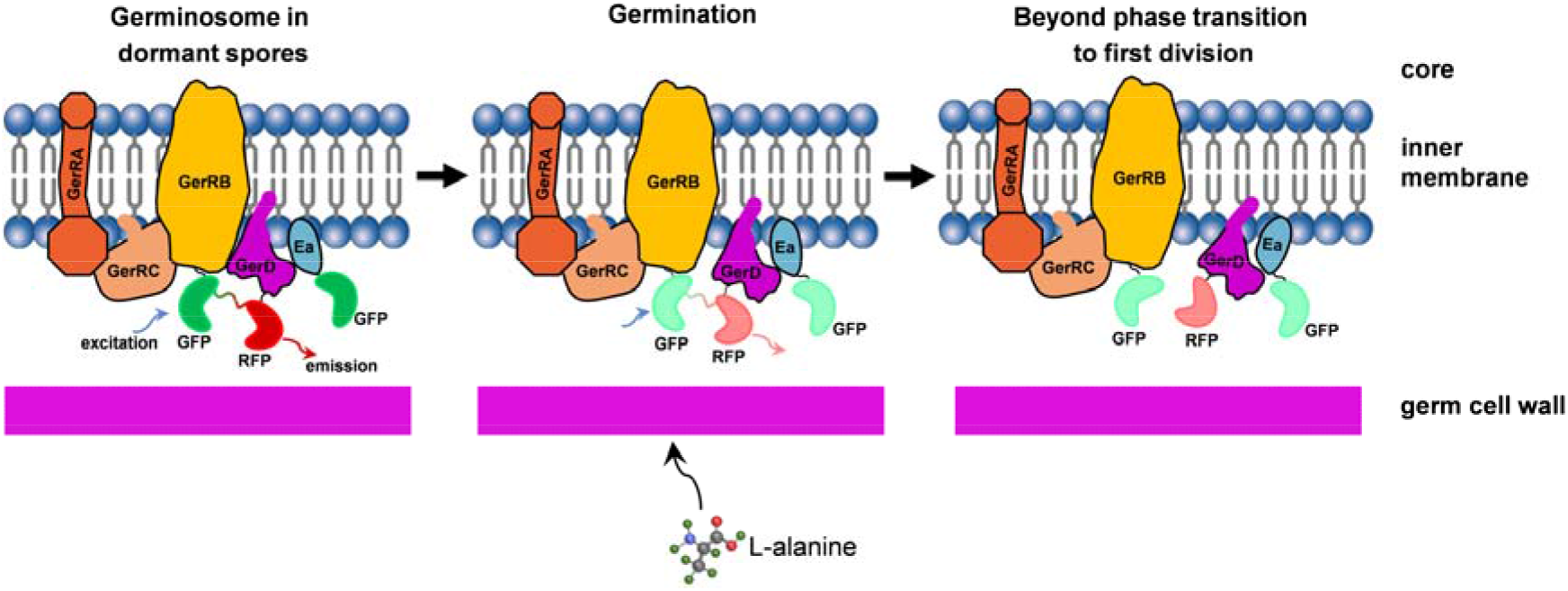
A proposed model of germinosome dynamics during germination triggered by L-alanine in *B. cereus* spores. The left most panel shows: i) FRET positive germinosome formation due to close interaction between GerR-SGFP2 and GerD-mScarlet-I; the darker green-red line between GFP (deep green) and RFP (deep red) indicates the energy transfer path in the FRET event between GerR and GerD; ii) likely colocalization between SpoVAEa and GerD proteins, albeit transient; GFP – SGFP2; RFP – mScarlet-I; Ea – SpoVAEa. The middle panel shows: i) FRET positive germinosomes may be lowered in intensity following the phase transition in germination initiation caused by L-alanine; the FRET signal (light green-red line) between GFP (light green) and RFP (light red) has become weak, indicating that close interaction between GerR and GerD has been gradually lost, consistent with GerD-mScarlet-I and GerRB-SGFP2 moving apart; ii) the SpoVAEa-SGFP2 and GerD-mScarlet-I fluorescence intensities have decreased upon initiation of germination. The right most panel shows: i) hypothesized loss of FRET positive germinosomes. Some GerR-SGFP2 foci (light green) and GerD-mScarlet-I foci (light red) may continue to exist, indicated by colocalization of GerR and GerD after the phase transition; ii) some GerD foci also continue to exist and likely colocalize, perhaps transiently with SpoVAEa.

In summary, the SpoVAEa-SGFP2 protein exhibits random movement on the outer surface of spores’ IM and a likely at least transient co-localization with GerD-mScarlet-I in dormant spores of *B. cereus;* this transient colocalization may be a means of transduction of a signal from a germinosome to SpoVA channels triggering CaDPA release from spores. This latter idea is certainly worth studying further. Studying spore germination by phase-contrast microscopy suggested that expression of GerR-SGFP2 or SpoVAEa-SGFP2 with GerD-mScarlet-I from a plasmid leads to more heterogeneity and lower efficiency of spore germination in *B. cereus*, pointing to the need for future studies to investigate the stoichiometry of the germinosome components in *B. cereus* in more detail. The dynamics of germination showed that germinosome foci composed of GerR-SGFP2 and GerD-mScarlet-I were lost soon after the phase transition. Further work related to the machinery of spore germination should likely focus on detailed studies of interactions between elements of the SpoVA channel, GerD and GR subunits.

## MATERIALS AND METHODS

### Recombinant plasmids and *B. cereus* strains

The recombinant plasmids and *B. cereus* strains used in this study are listed in Table 1. All primers used are listed in Table S1. The recombinant plasmids were constructed as described in previous studies (7, 20). Briefly, the region of 226 bp located in the upstream region of the *spoVA* operon was considered as the promoter region of *spoVAEa* gene and named PEa. The PEa fragment was inserted into pHT315 between *Kpn* I and *Xba* I sites resulting in plasmid pHT315-PEa. Next, the *spoVAEa* (BC_4065) gene was amplified from genomic DNA of *B. cereus* ATCC 14579 (GenBank: AE016877) using a pair of primers, 315_YW-42 and 315_YW-43. The *SGFP2* gene with stop codons was fused to the 3’ end of the *spoVAEa* gene using a two-fusion PCR. The fusion product was inserted into pHT315-PEa between *Xba* I and *Hind* III sites. The resulted ligation product was transformed into competent *E. coli* cells and selection of positive clones giving plasmid pHT315-f14. The fusion fragment *PD-gerD-mScarlet-I* was amplified from plasmid pHT315-f05 and inserted into pHT315-f14 between *Kpn* I and *EcoR* I sites, giving plasmid pHT315-f15. The correct construction of recombinant plasmids pHT315-f14 and pHT315-f15 was confirmed by sequencing, followed by electroporation into competent *B. cereus* ATCC 14579 cells, and selection and confirmation with colony PCR of an erythromycin-positive single colony.

**TABLE 1.**
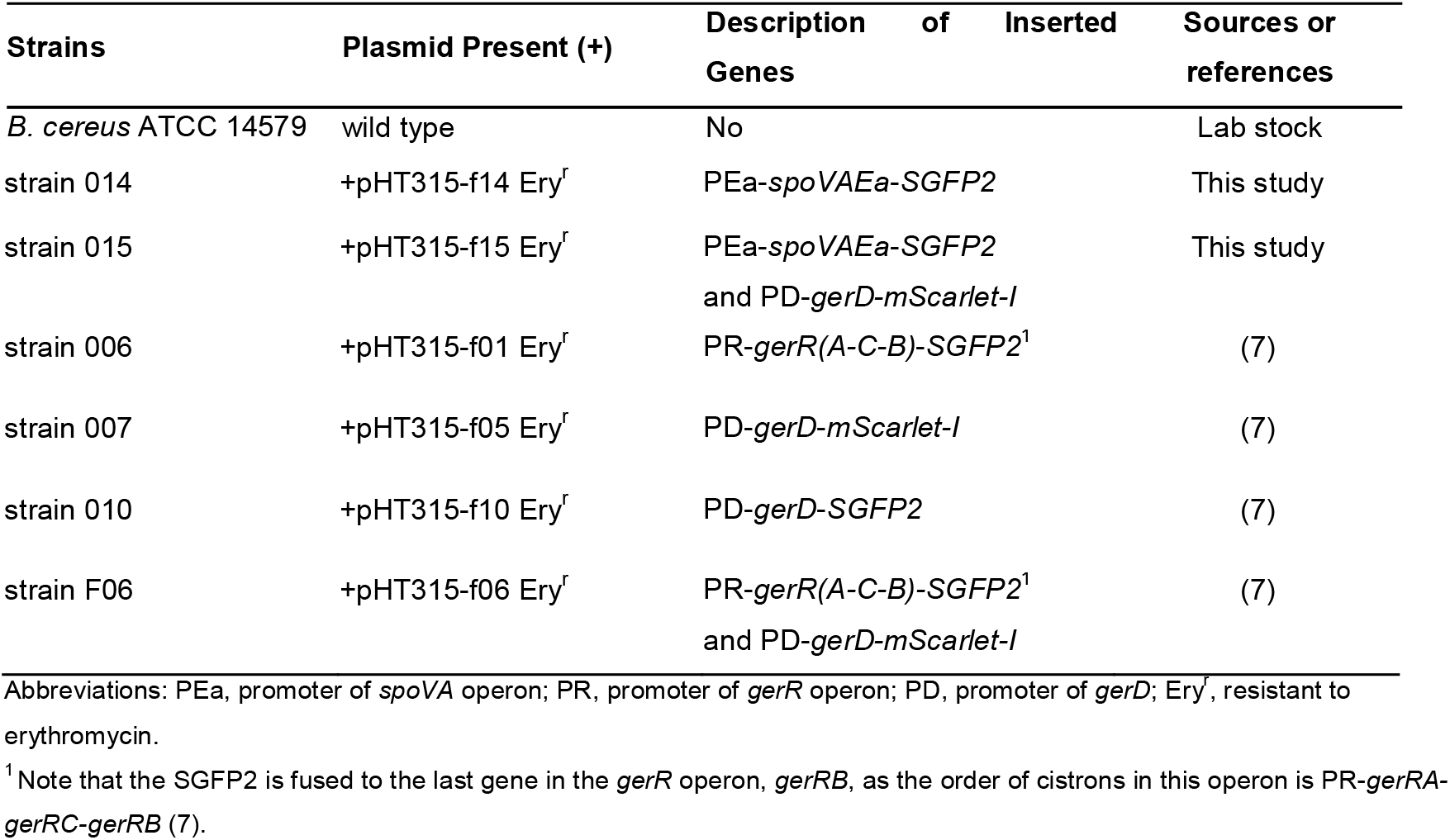
*B. cereus* strains and plasmids used in this study

### High frequency time-lapse image acquisition and analysis

Dormant spores of *B. cereus* strain 014 were prepared and purified as described in previous work (20). A Nikon Eclipse Ti-E microscope (Nikon Instruments, Tokyo, Japan) equipped with a sCmos camera (Hamamatsu Flash 4.0 V2, Hamamatsu City, Japan) and wide-field fluorescence components was used to capture 100 frames of 14-bit SGFP2 images (excitation at 488 nm and emission at 535 nm) with an exposure time of 50ms for each frame and no delay interval. An individual spore located in 100 frames was selected, duplicated, and analysed by the plugin Adrian’s FWHM in ImageJ. The percent change of FWHM in the second frame to the hundredth frame relative to the FWHM in the first frame was calculated, and the graph was made by the software of GraphPad Prism version 9.3.

### Images of SpoVAEa-SGFP2 expressed in spores of *B. cereus* strains 014 and 015; acquisition and analysis

The preparation of *B. cereus* dormant spores and their visualization were carried out as described in our previous work (20). Spores of *B. cereus* strain 014 expressing SpoVAEa-SGFP2 and spores of *B. cereus* strain 015 expressing SpoVAEa-SGFP2 and GerD-mScarlet-I were captured in the phase-contrast and SGFP2 (excitation at 470 nm and emission at 516 nm) channels using a Nikon Eclipse Ti-E microscope. Images were analysed by the ObjectJ SporeAnalyzer_1c.ojj in Fiji/ImageJ (https://sils.fnwi.uva.nl/bcb/objectj/examples/SporeAnalyzer/MD/SporeAnalyzer.html). Three independent experiments were performed, and the data were analyzed by the software of GraphPad Prism version 9.3.

### Colocalization assays and data analysis

Dormant spores of *B. cereus* strains 015 and 007 were prepared and purified as described in our previous work (20). Spores of *B. cereus* strains 015 and 007 were captured in three channels: phase-contrast, SGFP2 (excitation at 470 nm and emission at 516 nm) and mScarlet-I (excitation at 555 nm and emission at 593 nm) using a Nikon Eclipse Ti-E microscope. All acquired images in the colocalization assay were processed with ImageJ. The SGFP2 and mScarlet-I images were used to calculate the co-localization indicator Pearson’s coefficient by the plugin JACoP in ImageJ (22).

### Germination assays by time-lapse imaging and data processing

Dormant spores of *B. cereus* strains F06, 006, 015, 014, 007 and wild-type spores were prepared and purified as described previously (20). Microscope slides were prepared as described previously (30). Briefly, a 65 μl size Gene frame with 0.25 mm thickness (Thermofisher Scientific, The Netherlands, Cat. No.: AB0577) was attached on the center of a normal microscope slide. A liquid mixture for an agarose pad was made with a 1:1 mixture of 2× germination buffer (see below) and 2% agarose in a heat block at 55°C. 60 μl of the liquid mixture was pipetted on the area of frame, immediately pressed with another slide and placed at 4°C for at least 20 min to solidify.

Dormant spores suspended in ice-cold PBS (pH 7.4) were heat activated for 15 min at 70°C and washed three times with ice-cold PBS (pH 7.4) by centrifugation at 14,300 ×*g* for 15 min at 4°C. The heat-treated spores were suspended in ice-cold germination buffer (50 mM Tris-HCl (pH 7.4), 10 mM NaCl and 100 mM L-alanine) at an OD600 of 15. The spore suspension (1.3 μl) was dropped onto the solid agarose pad, immediately covered by a cover slip (18×18 mm) and was now ready for time-lapse microscopy.

A Nikon Eclipse Ti-E microscope (Nikon Instruments, Tokyo, Japan) equipped with an sCmos camera (Hamamatsu Flash 4.0 V2, Hamamatsu City, Japan), phase-contrast, and wide-field fluorescence components was used to track germination of *B. cereus* spores for 4 hours with 10 min intervals. Spores of *B. cereus* strain F06 expressing GerR-SGFP2 and GerD-mScarlet-I were captured by four images, phase-contrast, SGFP2 fluorescence (excitation at 470 nm and emission at 516 nm), mScarlet-I (excitation at 555 nm and emission at 593 nm) and FRET (excitation at 470 nm and emission at 593 nm). Spores of *B. cereus* strain 006 expressing GerR-SGFP2 and spores of *B. cereus* strain 014 expressing SpoVAEa-SGFP2 were captured by phase-contrast and SGFP2 images. Spores of *B. cereus* 015 expressing SpoVAEa-SGFP2 and GerD-mScarlet-I were captured by phase-contrast, SGFP2 and mScarlet-I images. Spores of *B. cereus* 007 expressing GerD-mScarlet-I were captured by phase-contrast and mScarlet-I images.

All 16-bit type images taken in germination assays were converted to 32-bit type. Selection and measurement of the area of background in samples without an image were carried out, and background was subtracted by the tool of Process—Math—Subtract in Fiji/ImageJ. The germinated spores were analysed and various intensities of individual spores measured using the ObjectJ SporeTrackerC_1h.ojj in Fiji/ImageJ (https://sils.fnwi.uva.nl/bcb/objectj/examples/sporetrackerc/MD/SporeTrackerC.html). Fluorescence intensities of plasmid-containing spores that were lower than that of wild-type spores were considered as aberrant values and were not counted. Three independent experiments were performed. The intensities of wild type spores were subtracted from the intensities of plasmid-containing, and the differences used to carry out statistical comparisons by the software GraphPad Prism version 9.3. Additionally, the calculation of NFRET (normalized FRET) was performed by our published method (7).

## Supporting information

Supplementary data SpoVAEa B. cereus

## ACKNOWLEDGMENTS

We acknowledge the Van Leeuwenhoek Center for Advanced Microscopy (LCAM) at the University of Amsterdam for offering the microscopy platform. We would like to thank Ronald M. P. Breedijk for his help in microscopy. We appreciate Juan Wen for sharing the raw data on the high frequency time-lapse images of SpoVA-SGFP2 in one *B. subtilis* spore. Finally, Yan Wang acknowledges the China Scholarship Council for her PhD scholarship.

We declare no conflicts of interest.

## References

1. Ehling-Schulz M, Lereclus D, Koehler TM. 2019. The *Bacillus cereus* Group: *Bacillus* species with pathogenic potential. Microbiol Spectr 7. doi:10.1128/microbiolspec.GPP3-0032-2018.

2. Jessberger N, Dietrich R, Granum PE, Märtlbauer E. 2020. The *Bacillus cereus* food infection as multifactorial process. Toxins (Basel) 12. doi:10.3390/toxins12110701.

3. Jovanovic J, Ornelis VFM, Madder A, Rajkovic A. 2021. *Bacillus cereus* food intoxication and toxicoinfection. Compr Rev Food Sci Food Saf 20:3719–3761. doi:10.1111/1541-4337.12785.

4. McKenney PT, Driks A, Eichenberger P. 2013. The *Bacillus subtilis* endospore: assembly and functions of the multilayered coat. Nat Rev Microbiol 11:33–44. doi:10.1038/nrmicro2921.

5. Pelczar PL, Setlow P. 2008. Localization of the germination protein GerD to the inner membrane in *Bacillus subtilis* spores. J Bacteriol 190:5635–5641. doi:10.1128/JB.00670-08.

6. Pelczar PL, Igarashi T, Setlow B, Setlow P. 2007. Role of GerD in germination of *Bacillus subtilis* spores. J Bacteriol 189:1090–1098. doi:10.1128/JB.01606-06.

7. Wang Y, Breedijk RMP, Hink MA, Bults L, Vischer NOE, Setlow P, Brul S. 2021. Dynamics of germinosome formation and FRET-based analysis of interactions between GerD and germinant receptor subunits in *Bacillus cereus* spores. Int J Mol Sci 22. doi:10.3390/ijms222011230.

8. Fort P, Errington J. 1985. Nucleotide sequence and complementation analysis of a polycistronic sporulation operon, *spoVA*, in *Bacillus subtilis*. J Gen Microbiol 131:1091–1105. doi:10.1099/00221287-131-5-1091.

9. Perez-Valdespino A, Li Y, Setlow B, Ghosh S, Pan D, Korza G, Feeherry FE, Doona CJ, Li Y-Q, Hao B, Setlow P. 2014. Function of the SpoVAEa and SpoVAF proteins of *Bacillus subtilis* spores. J Bacteriol 196:2077–2088. doi:10.1128/JB.01546-14.

10. Vepachedu VR, Setlow P. 2007. Analysis of interactions between nutrient germinant receptors and SpoVA proteins of *Bacillus subtilis* spores. FEMS Microbiol Lett 274:42–47. doi:10.1111/j.1574-6968.2007.00807.x.

11. Wen J, Vischer NO, de. Vos AL, Manders EMM, Setlow P, Brul S. 2021. Organization and dynamics of the SpoVAEa protein, and its surrounding inner membrane lipids upon germination of *Bacillus subtilis* spores. bioRxiv. doi:10.1101/2021.11.20.469378.

12. Moir A, Cooper G. 2015. Spore germination. Microbiol Spectr 3. doi:10.1128/microbiolspec.TBS-0014-2012.

13. Shen A, Edwards AN, Sarker MR, Paredes-Sabja D. 2019. Sporulation and germination in Clostridial pathogens. Microbiol Spectr 7. doi:10.1128/microbiolspec.GPP3-0017-2018.

14. Christie G, Setlow P. 2020. *Bacillus* spore germination: Knowns, unknowns and what we need to learn. Cell Signal 74:109729. doi:10.1016/j.cellsig.2020.109729.

15. Swerdlow BM, Setlow B, Setlow P. 1981. Levels of H+ and other monovalent cations in dormant and germinating spores of *Bacillus megaterium*. J Bacteriol 148:20–29. doi:10.1128/jb.148.1.20-29.1981.

16. Li Y, Davis A, Korza G, Zhang P, Li Y-Q, Setlow B, Setlow P, Hao B. 2012. Role of a SpoVA protein in dipicolinic acid uptake into developing spores of *Bacillus subtilis*. J Bacteriol 194:1875–1884. doi:10.1128/JB.00062-12.

17. Vary JC, Halvorson HO. 1965. Kinetics of germination of *Bacillus* spores. J Bacteriol 89:1340–1347. doi:10.1128/jb.89.5.1340-1347.1965.

18. Hashimoto T, Frieben WR, Conti SF. 1969. Microgermination of *Bacillus cereus* spores. J Bacteriol 100:1385–1392. doi:10.1128/jb.100.3.1385-1392.1969.

19. Kong L, Zhang P, Wang G, Yu J, Setlow P, Li Y-Q. 2011. Characterization of bacterial spore germination using phase-contrast and fluorescence microscopy, Raman spectroscopy and optical tweezers. Nat Protoc 6:625–639. doi:10.1038/nprot.2011.307.

20. Wang Y, Boer R de, Vischer N, van Haastrecht P, Setlow P, Brul S. 2020. Visualization of germination proteins in putative *Bacillus cereus* germinosomes. Int J Mol Sci 21. doi:10.3390/ijms21155198.

21. Hornstra LM, Vries YP de, Vos WM de, Abee T, Wells-Bennik MHJ. 2005. *gerR*, a novel *ger* operon involved in L-alanine-and inosine-initiated germination of *Bacillus cereus* ATCC 14579. Appl Environ Microbiol 71:774–781. doi:10.1128/AEM.71.2.774-781.2005.

22. Bolte S, Cordelières FP. 2006. A guided tour into subcellular colocalization analysis in light microscopy. J Microsc 224:213–232. doi:10.1111/j.1365-2818.2006.01706.x.

23. Adler J, Parmryd I. 2010. Quantifying colocalization by correlation: the Pearson correlation coefficient is superior to the Mander’s overlap coefficient. Cytometry A 77:733–742. doi:10.1002/cyto.a.20896.

24. Cabrera-Martinez R-M, Tovar-Rojo F, Vepachedu VR, Setlow P. 2003. Effects of overexpression of nutrient receptors on germination of spores of *Bacillus subtilis*. J Bacteriol 185:2457–2464. doi:10.1128/JB.185.8.2457-2464.2003.

25. Gao XW. 2022. *Bacillus cereus* spore and cell proteome dynamics. PhD thesis. Swammerdam Institute for Life Sciences, Amsterdam, The Netherlands.

26. Hornstra LM, Vries YP de, Wells-Bennik MHJ, Vos WM de, Abee T. 2006. Characterization of germination receptors of *Bacillus cereus* ATCC 14579. Appl Environ Microbiol 72:44–53. doi:10.1128/AEM.72.1.44-53.2006.

27. Zhang J, Griffiths KK, Cowan A, Setlow P, Yu J. 2013. Expression level of *Bacillus subtilis* germinant receptors determines the average rate but not the heterogeneity of spore germination. J Bacteriol 195:1735–1740. doi:10.1128/JB.02212-12.

28. Griffiths KK, Zhang J, Cowan AE, Yu J, Setlow P. 2011. Germination proteins in the inner membrane of dormant Bacillus subtilis spores colocalize in a discrete cluster. Mol Microbiol 81:1061–1077. doi:10.1111/j.1365-2958.2011.07753.x.

29. Artzi L, Alon A, Brock KP, Green AG, Tam A, Ramírez-Guadiana FH, Marks D, Kruse A, Rudner DZ. 2021. Dormant spores sense amino acids through the B subunits of their germination receptors. Nat Commun 12:6842. doi:10.1038/s41467-021-27235-2.

30. Pandey R, Beek A ter, Vischer NOE, Smelt JPPM, Brul S, Manders EMM. 2013. Live cell imaging of germination and outgrowth of individual *Bacillus subtilis* spores; the effect of heat stress quantitatively analyzed with SporeTracker. PLoS One 8:e58972. doi:10.1371/journal.pone.0058972.

