## Supplementary data SpoVAEa B. cereus for "Visualization of SpoVAEa protein dynamics in dormant spores of *Bacillus cereus* and dynamic changes in their germinosomes and SpoVAEa during germination"

**Supplementary Materials**

**TABLE S1**. Primers used in this study

| Primers | Sequence (5'-3') | Purposes or functions |
| --- | --- | --- |
| 315_YW-42 | GGGGTACCAGGTGTTTCATAATGAGTGG | *Kpn* I-*spoVA* promoter-Fw |
| 315_YW-43 | GCTCTAGACTTTCATTCACCCCTTCAC | *Eco*R I-*spoVA* promoter-Rv |
| 315_YW-44 | GCTCTAGATTGGTCACAGATGGAGAT | *Xba* I-*spoVAEa*-Fw |
| 315_YW-45 | TCCTCGCCCTTGCTCACCATgctgccgctgccgctgcc  GGCTTCGTCATAAAAACCA | *spoVAEa-(GS)_3_-SGFP2-*Rv |
| 315_YW-46 | TGGTTTTTATGACGAAGCCggcagcggcagcggcagc  ATGGTGAGCAAGGGCGAGGA | *spoVAEa-(GS)_3_-SGFP2-*Fw |
| 315_YW13 | CCCAAGCTTTTACTTGTACAGCTCGTCCAT | *SGFP2*‐*Hin*d III‐Rv |
| 315_YW-47 | CAAATGGGGAAGTCTCGT | *spoVAEa-*sequencing-Rv |

a. Restriction cleavage sites are underlined.

b. The six-residue flexible linker (Gly-Ser-Gly-Ser-Gly-Ser) coding sequence is written in lower‐case letters.

**TABLE S2**. The numbers of germinated spores of different *B. cereus* strains examined in three independent experiments

| *B. cereus* strains | replication | 1min | 10min | >20min | total |
| --- | --- | --- | --- | --- | --- |
| strain F06 | replication 1 | 74 | 10 | 5 | 89 |
|  | replication 2 | 87 | 12 | 9 | 108 |
|  | replication 3 | 56 | 9 | 5 | 70 |
| strain 006 | replication 1 | 39 | 7 | 4 | 50 |
|  | replication 2 | 43 | 4 | 3 | 50 |
|  | replication 3 | 78 | 10 | 4 | 92 |
| strain 015 | replication 1 | 13 | 9 | 23 | 45 |
|  | replication 2 | 69 | 18 | 25 | 122 |
|  | replication 3 | 48 | 13 | 19 | 80 |
| strain 014 | replication 1 | 35 | no | no | 35 |
|  | replication 2 | 228 | 2 | 3 | 233 |
|  | replication 3 | 62 | 1 | 2 | 65 |
| strain 007 | replication 1 | 29 | 3 | 6 | 38 |
|  | replication 2 | 82 | 13 | 12 | 107 |
|  | replication 3 | 59 | 8 | 16 | 83 |
| wild type | replication 1 | 29 | no | no | 29 |
|  | replication 2 | 23 | no | no | 23 |
|  | replication 3 | 263 | 2 | no | 265 |

**TABLE S3**. Statistical comparisons between each timepoint and 0 min of intensities in Fig. 6 of channels from germinating spores of *B. cereus* strain F06

| Groups | Comparisons  between times^1^ | Channels | | | |
| --- | --- | --- | --- | --- | --- |
|  |  | PH3 | SGFP2 | mScarlet-I | FRET |
| germX_1 | 10 min | ns | ns | ns | ns |
|  | 20 min | **** | ns | ns | * |
|  | 30 min | **** | ns | ns | *** |
|  | 40 min | **** | ns | *** | *** |
|  | 50 min | **** | ns | *** | *** |
|  | 60 min | **** | ns | *** | **** |
| germX_10 | 10 min | ns | ns | ns | ns |
|  | 20 min | ns | ns | ns | ns |
|  | 30 min | **** | ns | ns | ns |
|  | 40 min | **** | ns | * | * |
|  | 50 min | **** | ns | ** | ** |
|  | 60 min | **** | ns | ** | ** |

ns, not significant; *, *P* < 0.05; **, *P* < 0.01, ***, *P* < 0.001; ****, *P* < 0.0001

^1^Comparisons were between intensities at indicated times and the time above, except for the 10 min samples that were compared to spores at time 0.

**TABLE S4**. Statistical comparison of vertical pairs’ intensities in Fig. S2 of channels from germinating spores of *B. cereus* strains 006, 007 and 014.

| Groups | Comparisons between times^1^ | strain 006 | strain 007 | strain 014 |
| --- | --- | --- | --- | --- |
|  |  | SGFP2 channel | mScarlet-I channel | SGFP2 channel |
| gerX_1 | 10 min | ns | ns | ** |
|  | 20 min | ** | *** | *** |
|  | 30 min | *** | **** | ** |
|  | 40 min | **** | **** | * |
|  | 50 min | **** | **** | ns |
|  | 60 min | **** | **** | ns |
| germX_10 | 10 min | ns | ns | ns |
|  | 20 min | * | ns | ns |
|  | 30 min | ns | ** | ns |
|  | 40 min | ns | **** | ns |
|  | 50 min | ns | **** | ns |
|  | 60 min | ns | **** | ns |

ns, not significant; *, *P* < 0.05; ***, *P* < 0.001; ****, *P* < 0.0001.

^1^Comparisons were between intensities at indicated times and the time above, except for the 10 min samples that were compared to spores at time 0.

**TABLE S5**. Statistical comparisons of vertical pairs of intensities in Fig. 7 of channels from germinating spores of *B. cereus* strain 015

| Groups | Comparisons between times^1^ | Channels | | |
| --- | --- | --- | --- | --- |
|  |  | PH3 | SGFP2 | mScarlet-I |
| germX_1 | 10 min | **** | ns | ns |
|  | 20 min | **** | ns | ns |
|  | 30 min | **** | ns | *** |
|  | 40 min | **** | ns | **** |
|  | 50 min | **** | ns | **** |
|  | 60 min | **** | ns | **** |
| germX_10 | 10 min | ns | ns | ns |
|  | 20 min | **** | ns | ns |
|  | 30 min | **** | ns | * |
|  | 40 min | **** | ns | ** |
|  | 50 min | **** | ns | **** |
|  | 60 min | **** | ns | **** |

ns, not significant; ****, *P* < 0.0001.

^1^Comparisons were between intensities at indicated times and the time above, except for the 10 min samples that were compared to spores at time 0.

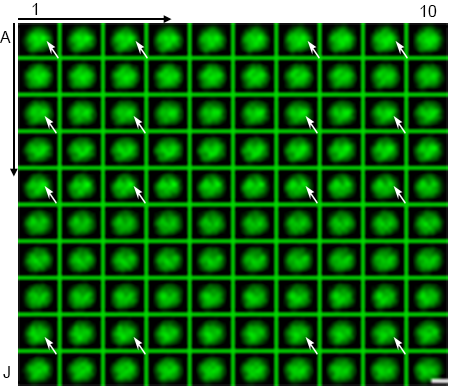

**FIG S1**. A montage of 100 frames (A1 to J10) of *B. cereus* spore 2 in Fig. 1. Each frame was acquired with 488 nm excitation light and at 535 nm emission light with an exposure time of 50ms and no delay interval. The scale bar is 1 µm.

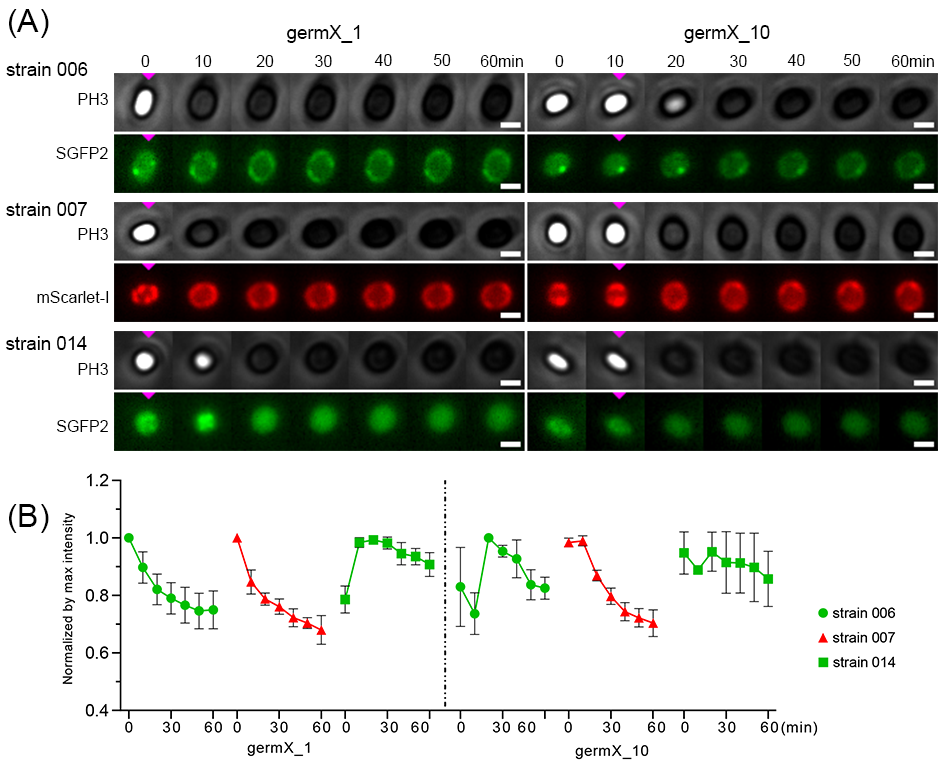

**FIG S2**. Dynamic changes in germinated spores of *B. cereus* strains 006, 007, and 014. Panel A, visualization of dynamics of two representative germinated spores of each strain at 10 min intervals over 60 min. Channels PH3 (phase contrast) and SGFP2 in the top layer are the groups germX_1 and germX_10 in spores of *B. cereus* strain 006. Channels PH3 and mScarlet-I in the middle layer are the groups germX_1 and germX_10 in spores of *B. cereus* strain 007. Channels PH3 and SGFP2 in the bottom layer are the groups germX_1 and germX_10 in spores of *B. cereus* strain 014. PH3, phase contrast. The pink triangles indicate the initiation of spore germination. The scale bar is 1 µm. Panel B, the line charts of SGFP2 (green) or mScarlet-I (red) channel intensities. Left column: group germX_1; right column: group germX_10. Data are shown as the mean with SD from three independent experiments. The numbers of analyzed germinated spores of *B. cereus* strains 006, 007 and 014 are given in Table S2. The statistical analyses of results are given in Table S4.
